# A tool for federated training of segmentation models on whole slide images

**DOI:** 10.1101/2021.08.17.456679

**Authors:** Brendon Lutnick, David Manthey, Jan U. Becker, Jonathan E. Zuckerman, Luis Rodrigues, Kuang Yu. Jen, Pinaki Sarder

## Abstract

The largest bottleneck to the development of convolutional neural network (CNN) models in the computational pathology domain is the collection and curation of diverse training datasets. Training CNNs requires large cohorts of image data, and model generalizability is dependent on training data heterogeneity. Including data from multiple centers enhances the generalizability of CNN based models, but this is hindered by the logistical challenges of sharing medical data. In this paper we explore the feasibility of training our recently developed cloud-based segmentation tool (Histo-Cloud) using federated learning. We show that a federated trained model to segment interstitial fibrosis and tubular atrophy (IFTA) using datasets from three institutions is comparable to a model trained by pooling the data on one server when tested on a fourth (holdout) institution’s data. Further, training a model to segment glomeruli for a federated dataset (split by staining) demonstrates similar performance.

## Introduction

As the practice of digitizing histological slides has become common practice^1^, the field of computational pathology has exploded. Modern image analysis technologies (such as deep learning^2^) are increasingly being applied to examine digitized whole slide images (WSIs). The maturation of convolutional neural networks (CNNs)^3^ (a specialized subset of deep learning) for the analysis and segmentation of natural images has led to widespread adoption of this technology in the field of computational pathology. CNNs have shown promising results for state of the art computational pathology image analysis tasks including tissue segmentation^4–8^, disease classification^9–13^, and outcome prediction^14,15^. Training these networks is enhanced by access to diverse WSI datasets, as greater data variability is known to enhance model robustness^16^. For histological tissue, stained and scanned digitally as WSIs, the institution where data is prepared often has a large effect on the quality and appearance of the tissue^17^. Institution specific factors such as tissue preparation and staining protocol, as well as any demographic biases can have a large effect on the resulting WSIs. Practically this means gathering training data from multiple institutions. However sharing medical data across institutions can be complicated by regulatory challenges^18^, limiting the scope of collaboration and therefore the generalizability of computational pathology tools.

Federated learning was recently proposed as an efficient solution for decentralized training of models without sharing data^19,20^. Training a network on a federated dataset uses multiple rounds of local training performed on hardware located at the data source, the learned network parameters are shared and averaged between each round to avoid divergence between training sites. At the core of federated learning is federated averaging (FedAvg)^21^, which is simply a weighted average of the network weights across training sites, performed at pre-selected intervals (Fig. 1). FedAvg has been practically shown to achieve convergence with proper hyperparameter tuning^22^. Originally proposed for smartphone natural language processing tasks where data sharing is limited by a limited network bandwidth and privacy concerns, federated learning has recently gained the interest of computational researchers in the medical field^23^. Computational pathology datasets are a perfect candidate for federated learning where both file sizes of WSIs (gigapixels) and regulatory limitations hinder data sharing.

**Fig. 1.**
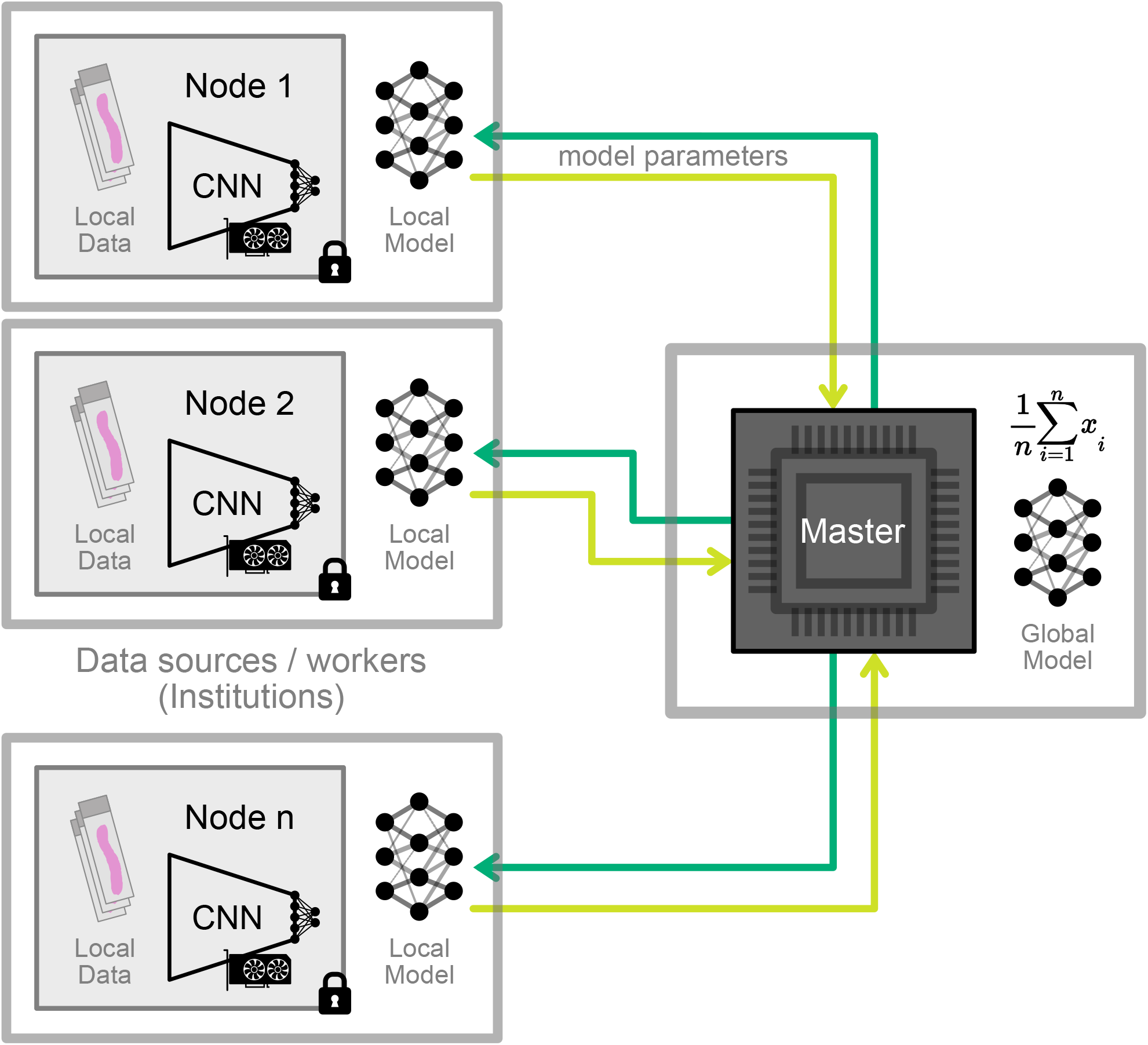
The federated learning schematic. A schematic example of federated learning. Multiple worker nodes store data and model parameters locally at the institution of origin. The data stored on these worker nodes is never shared, and the nodes perform local training using this data upon the request of the master server. The local models are then shared with the master server who performs parameter averaging, before sending the updated global model back to the worker nodes for further local training. This process is repeated iteratively throughout the training process, until model convergence.

To show the feasibility of federated learning on pathology data in the real world, we have created a pipeline for federated segmentation on WSIs capable of deployment across multiple institutions. This pipeline is deployed in the cloud for easy access for data viewing and annotation by each site’s respective constituents. This is a companion work to our recently published Histo-Cloud segmentation tool^8^; it shows the feasibility for training Histo-Cloud in a federated setup. Histo-Cloud is a cloud-based tool for segmentation of WSIs. It combines the digital slide archive (DSA)^24^ for WSI data management, HistomicsUI for WSI viewing and annotation, and a modified version of the DeepLab V3+ network^5^ for WSI segmentation.

## Results

Federated training was performed on data distributed across three discreate servers (workers). A fourth server acted as the master server, performing parameter averaging, and training synchronization, a schematic is available in Fig. 1. In each server, data is stored on an instance of the DSA^24^, and our Histo-Cloud plugin^8^ is responsible for network training. This plugin is capable of utilizing hardware acceleration for training, and uses two available GPUs in all three host machines for a total of 6 GPUs. We demonstrate the feasibility of federated segmentation of WSIs with two case studies:

### 1- Federated IFTA segmentation (divided by institution)

For the first case study interstitial fibrosis and tubular atrophy (IFTA) was segmented from WSIs from biopsies containing renal allograft nephropathy stained using periodic acid schiff (PAS). Three pathologists from different institutions each provided a minimum of 20 PAS stained WSIs. The WSIs per set were uniformly chosen from four IFTA classes defined based on ci/ct scores (0, 1, 2, & 3); ci/ct scoring is a method defined in Banff 2018 criteria^25^ for assessing transplant biopsies. A minimum of five slides per class were used for each set. The cases were reviewed to ensure the following selection criteria were met: (1) the amount of early or evolving IFTA with variable intermixed edema was minimized, (2) no active inflammation, (3) no prior history of rejection, and (4) cases were selected to represent the full range of IFTA severity. All types of IFTA, including classic, endocrinization, and thyroidization types, were included in the analysis, without distinguishing between the types. In total the pathologists from institutions 1-3 provided 20, 48, and 22 slides respectively. A holdout dataset was randomly selected by pooling 1/3^rd^ of the slides from each institution (29 slides total). We trained 5 models using this dataset: The first model was trained across three federated servers, training data for this study was split by institution of origin. For a baseline performance, a second model was trained centrally by pooling all the training data on a single server and using traditional gradient decent. Finally, to compare the performance in a data restricted setting, three additional models were trained using data from a single institution alone.

We note that IFTA boundaries are poorly defined, and subject to disagreement between pathologists^26^, receiver operating characteristic (ROC) curves were used to better capture the performance characteristics of our trained models. These were generated by applying a varying threshold to the network logits for the prediction of IFTA regions. To measure performance, we calculate the area under the curve (AUC) which is a common metric for measuring performance when a ROC curve is available.

Testing these models on the holdout set, we observed that central training and federated training of the IFTA model performed similarly both with *AUC = 0.95*. Performance fell when testing the models trained using a single institutions data, giving *AUC = 0.92, 0.87, & 0.91* respectively. ROC plots of the performance of the five models is highlighted in Fig. 2a. An example of IFTA segmentation on a holdout slide using the federated model is shown in Fig. 2c. Here we use the network logits to display the predictions as a probabilistic heatmap which we believe is better for the display of structures with poorly defined boundaries such as IFTA.

**Fig. 2.**
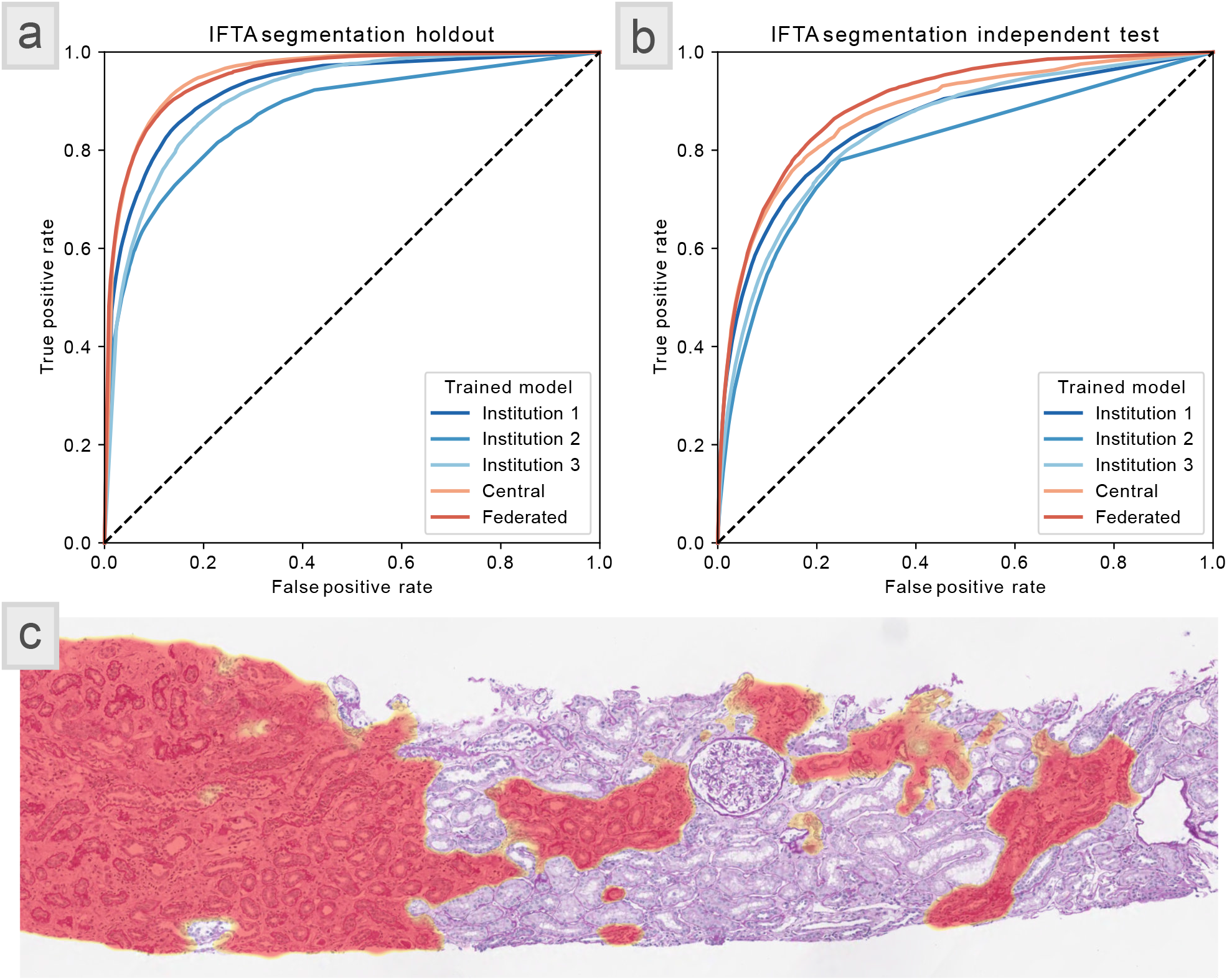
Multi institute IFTA segmentation performance, data split by institution. The segmentation performance of the trained IFTA models. This data was split by institution across three servers for federated training. Due to the subjective nature of IFTAboundaries, we use ROC curves and AUC to measure the segmentation performance. **[a]** ROC curves showing each models performance on a dataset of 29 holdout WSIs which were randomly selected from the same data as the training set. We observed that central training and federated training of the IFTA model performed similarly both with *AUC = 0.95*. Performance fell when testing the models trained using a single institutions data, giving *AUC = 0.92, 0.87, & 0.91* respectively. **[b]** ROC curves showing each models performance on an independent test set of data containing 17 WSIs. This dataset was from an institution which did not provide any training data, and was annotated by an independent pathologist. Similar to the holdout set, the central and federated models outperformed the models trained on a single institution’s data. Interestingly the federated model performed best with *AUC = 0.90* and the central model also performed well with *AUC = 0.88*.The institutions 1, 2, & 3 had *AUC = 0.85, 0.81, & 0.84* respectively. **[c]** an example of IFTA segmentation using the federated model on a slide from the holdout dataset. The prediction of IFTA is shown here using a heatmap, which reflects the network confidence in IFTA segmentation.

A fourth pathologist from a different institution provided an additional 17 slides to be used as an independent testing dataset. When we applied the trained IFTA models to this independent set we observed a similar trend as the holdout set. Here the federated model performed best with *AUC = 0.90* and the central model also performed well with *AUC = 0.88*. Like the holdout set, performance of the models trained on a single institution was lower than federated or central models, with *AUC = 0.85, 0.81, & 0.84* respectively. ROC plots of the performance of the five models is highlighted in Fig. 2b.

### 2- Federated glomeruli segmentation (divided by stain)

As a further test of our method, we designed a second study focused on glomerular segmentation, with a goal of studying the effects of tissue staining on federated training. A training set of 75 human renal WSIs from transplant biopsies, stained with 25 periodic acid schiff (PAS), 25 hematoxylin & eosin (H&E), and 25 Masson’s trichome (TRI). These slides were selected from a single institution and ground truth glomeruli annotation was performed by a single annotator. Slides were divided by stain and uploaded to the three training servers (25 slides per server). Like the IFTA study, five models were trained: A federated model, a central model using all the data, and three models using each stain individually. A holdout set of 30 slides (10 from each stain) was selected from the same dataset as the training data.

Unlike IFTA, glomerular boundaries are well defined and are best displayed by directly using the network predictions. We convert these predictions to contours for display on the slide. Fig. 3c shows an example of glomerular boundaries predicted using the federated model on holdout slides of various stains. We choose to use Matthews correlation coefficient (MCC) to measure the performance of glomerular segmentation, as it is commonly used for binary segmentation tasks. On the holdout data the federated model and the central model had identical performance, both achieving *MCC = 0.91*, outperforming all the models trained using only one stain (*MCC = 0.88* H&E, *0.56* PAS, and *0.85* TRI). A violin plot of the holdout performance as a function of the model used is shown in Fig. 3a.

**Fig. 3.**
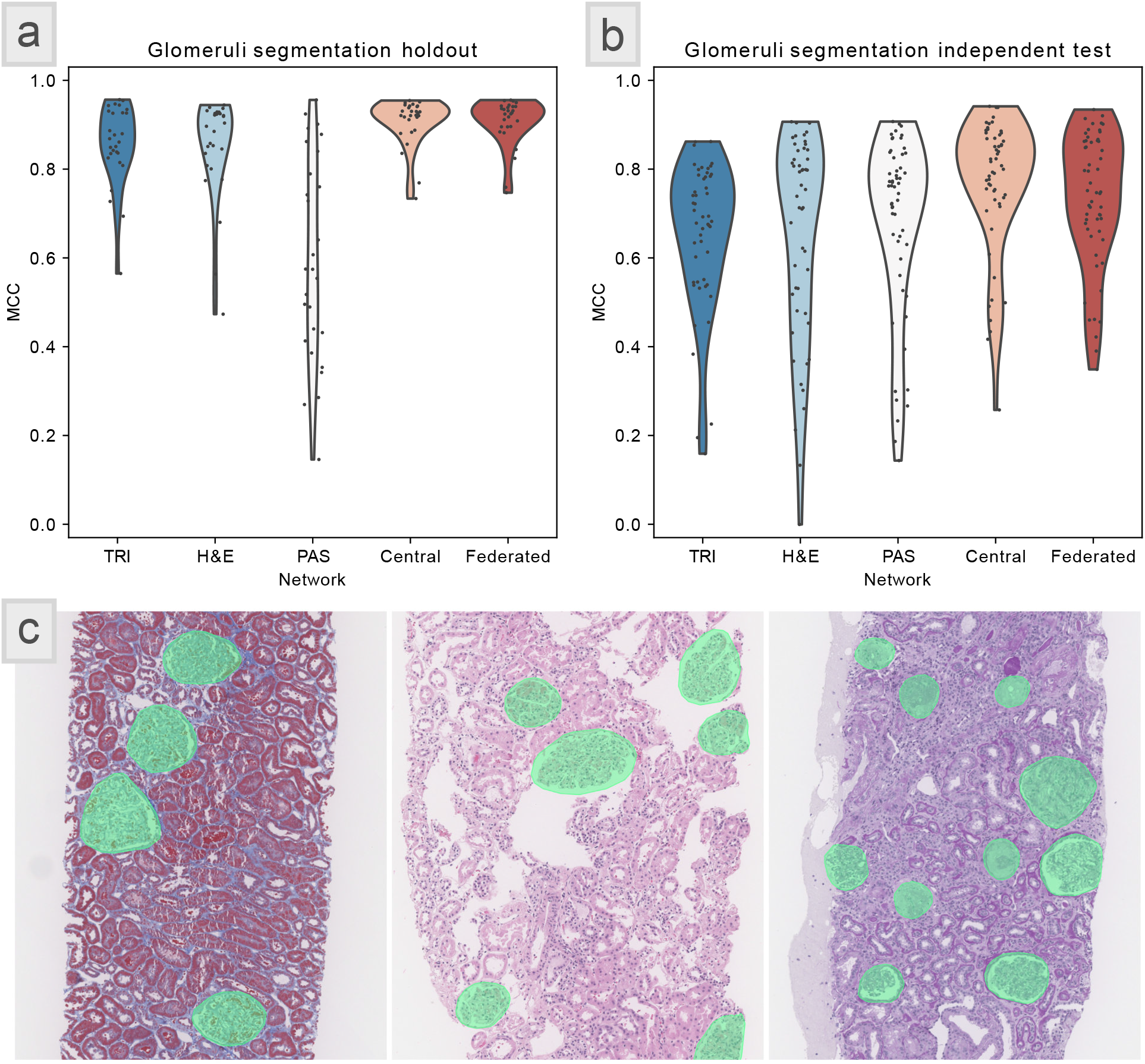
Glomeruli segmentation performance, data split by stain. The segmentation performance of the models trained for glomeruli segmentation. This data was split by stain across three servers for federated training. Because glomeruli have well defined boundaries we use MCCto calculate the segmentation performance, without varying the network prediction thresholds. **[a]** a violin plot showing the performance of each model on a dataset of 30 holdout WSIs which were randomly selected from the same data as the training set. We observed that central training and federated training of the models performed similarly both with MCC = 0.91. Performance fell when testing the models trained using a single stain, giving MCC = 0.85, 0.88, & 0.56 for TRI, H&E, & PAS stains respectively. **[b]** a violin plot showing the performance of each model on a dataset of 58 holdout WSIs which were selected from an independent institution. This dataset also included WSIs stained with Jones, which was not used for training. The federated model (MCC = 0.80) was outperformed by the central model (MCC = 0.83). However, the federated model still outperformed the models trained using a single stain alone (MCC =0.65 TRI, 0.69 H&E, & 0.78 PAS). **[c]** examples of glomeruli segmentation using the federated model on three slides from the holdout dataset. From left to right the slides are stained with trichrome, H&E, and PAS.

To further show the generalizability of the model, an independent testing set of 58 slides was chosen from a separate institution, and annotated by a separate annotator. This data included PAS, H&E, and TRI stains, as well as Jones stain. As expected, the segmentation performance was reduced on the independent test set. The federated model (*MCC = 0.80*) was outperformed by the central model (*MCC = 0.83*). However, the federated model still outperformed the models trained using a single stain alone (*MCC = 0.69* H&E, *0.78* PAS, and *0.65* TRI). A violin plot of the holdout performance as a function of the model used is shown in Fig. 3b.

## Discussion

Numerous examples of federated learning on medical data exist^27–30^, however at the time of writing computational pathology research on federated learning using WSIs is limited to a paper by Lu et al.^31^. Lu at al. trained a weakly supervised, multi instance learning model for subtyping breast cancer and renal cell carcinoma and predicting survival, while exploring the effects of differential privacy^32^ on model performance. Setting aside the complexities of network hyperparameter tuning, we argue that at its core, federated learning is a data organization and synchronization problem. While current applications in the literature describe the hyperparameters used for training, their data management and synchronization strategies lack details. Often federated learning research is performed locally in one machine, relying on simulated data sites^19^. For example, the details of the federated setup used by Lu et al.^31^ are not well described, and it is unclear if the federated training was simulated or actually performed across physically distinct servers. While results of simulation are valid for method development, we argue that the complexities of managing data and coordination across multiple training sites are a large logistical hurdle for real world applications of federated learning.

Our experiments (IFTA & glomeruli segmentation) show that not only does federating training for WSI segmentation converge, but the resultant model outperforms training done with a single dataset (institution orstain). Furthermore,the federated model performs on parwith a model trained traditionally with multiple datasets gathered at a central location. Most importantly, these experiments demonstrate the feasibility of training and coordinating federated segmentation models, managing datasets distributed across physically separate servers, and training in reasonable time.

We are not the first to propose federated segmentation, *L. Yi et al*. proposed SU-Net^33^ a federated network for brain tumor segmentation, which performed similarly to DeepLab^5^ for non-federated training. The first to train the DeepLab V3+^5^ architecture in a federated setup was *U. Michieli et al*.^34^ who used the VOC2012 dataset^35^ and simulated federated training on a single machine. In contrast, our federated approach offers comparable performance (both in training time and segmentation accuracy) to traditional training of DeepLab on gigapixel sized medical images (WSIs). Working efficiently with WSIs using CNNs requires a substantial amount of engineering effort, and the backbone of our code used fortraining was custom built to extract and process regions of interest from WSIs efficiently. We believe the ability to easily manage and annotate WSI data at each federated site using the DSA^24^ greatly enhances the real world applications of our method.

Throughout our training process, the newest segmentation model is available for testing at each data site, and could theoretically be used in a human in the loop approach to aid in the annotation of new WSIs similar to our previously described H-AI-L approach^7^. Newly added WSIs will automatically be incorporated into the training set at the beginning of each round of training.

Approaches such as peer-to-peer federated learning^36^ and swarm learning^37^ offer data synchronization strategies that do not require a central coordinating (master) server. While the lack of centralized training coordination may be beneficial for some tasks, we argue that for federated medical image segmentation, it is likely that only one group will be responsible for model development. Therefore, the ability to control and monitor training as well as adjust hyperparameters on one master server is ideal. The typical setup of federated learning in a medical setting will involve orders of magnitude less training sites than a task such as speech recognition, which has millions of potential training sites such as mobile phones. Multi-institute federated studies using medical images will require careful central coordination with recruitment and opt in by participating sites. The hardware for performing the training (at least for medical image analysis) is specialized, requiring IT setup and support at each institution. Our Histo-Cloud tool (which is used for training) is easy to setup making it ideal for this use case.

## Methods

An open source version of our code will be released with the publication of this work in a peer reviewed journal.

### Segmentation plugin

This work is heavily based upon our previously published Histo-Cloud tool^8^, where we modified the DeepLab V3+ architecture^5^ to work natively on WSIs and developed a series of plugins for running segmentation training and prediction in the cloud. This work was based on the Digital Slide Archive (DSA)^24^ an open source slide viewer and repository developed by Kitware Inc. Specifically these plugins were developed for accessibility in HistomicsUI, the slide viewing component of the DSA. Functions of the DSA can be controlled using a REST web API^38^, this includes the ability to trigger jobs by running the installed plugins as well as upload and download data stored in the DSA. Achieving federated learning using this system was straightforward. We use the requests python library^39^ to send REST calls to the federated workers, which all have the DSA and Histo-Cloud installed and are hosting the respective training datasets.

### Server coordination

In this pipeline, each site / institution has a worker node server with the DSA installed where training data is uploaded and annotated. Using the DSA data permissions can be set so that only eligible users from the institution have access to this data. A central (master) server manages the training cycle, uploading the global model parameters to each worker via the DSA REST API before requesting each run local training. Training Jobs are submitted to each worker (training site) using the Histo-Cloud training plugin^8^. The training job scheduling is handled by the DSA internally using *slicer_cli_web*, which uses Celery^40^ for task queue management, and RabbitMQ^41^ as a message broker. The job status is monitored by the master server until completion. Upon job completion the master server requests and downloads the resultant saved local model parameters from each worker node. These parameters are averaged by the master server and the global model is updated accordingly. The next round of training is then initiated: the global model is uploaded to each worker and is trained further before being downloaded and averaged. If training fails on one of the participating workers, then it is excluded from the rest of the training round, but participates in future training rounds.

### Data management

The training WSI data is uploaded to the DSA worker servers, where it was annotated by expert pathologists. Training data is placed in a folder created on each worker for easy access by the Histo-Cloud training plugin. A separate folder was created for the models produced by training and uploaded after federated averaging. The ID of these folders is known by the master server so it can submit training jobs specifying the data and models to be used for training.

### Training process

Training rounds involve parameter upload, training, parameter download, and federated parameter averaging across the worker and master nodes. The following pseudocode describes the training process:

**INIT** global model to ImageNet parameters
**WHILE** global training steps is less than total steps
**FOR** each client, **in parallel do**
**UPLOAD** global model parameters to clients
**CALL** TrainNetwork plugin **involving**
**INIT** network parameters with global model parameters
**TRAIN** for number of steps in round
**UPDATE** global training steps
**DOWNLOAD** trained local model
**END FOR**
**COMPUTE** average model parameters (FedAvg)
**SET** global model to FedAvg parameters
**END WHILE**

### Training setup

For training we used three physically distinct Linux servers running Ubuntu 18.04.5 LTS, with the DSA installed. The three servers had different hardware configurations, notably the graphics processing units (GPUs) were different across the servers. All computers had 2 GPUs that were produced by the Nvidia corporation and included:

1. Titan X Pascale (12GB VRAM) & GeForce GTX 1080 (8GB VRAM) - batch size 4
2. GeForce RTX 2080 Ti (11GB VRAM) & GeForce GTX 1080 (8GB VRAM) - batch size 4
3. 2X Quadro RTX 5000 (16 GB VRAM) - batch size 12

For training we used both available GPUs on each server and adjusted the batch size for each server to accommodate the individual VRAM (GPU memory) capacity of each.

### Training hyperparameters

The goal of federated averaging is to speed up training by removing the overhead of frequent communication between training sites. This is done by training locally for multiple steps before updating the central model parameters using FedAvg. Practically when optimizing the hyperparameters of our training loop, we found that using 1000 training steps between FedAvg achieved repeatable convergence. We trained for a total of 40 rounds (40,000 steps), using the momentum optimizer^42^ with an initial learning rate of 7e^-3^, using polynomial decay with a learning power of 0.9 and final learning rate of 0. To achieve stability at the start of training, we set the learning rate to 1e^-4^ for the first 750 steps. Finally, the gradients on the last layers of the network were scaled up by a factor of 10 to achieve faster convergence. These layers included the ASPP pooling layers and the layers in the decoder as defined by the DeepLab network architecture^5^.

All the models were trained using transfer learning with parameters inherited from a model pre-trained on the ImageNet dataset. Due to the low maximum batch size of 4 on two of the training servers, we did not train the batch normalization parameters. Training patches with a size of 512×512 pixels were extracted from the training WSIs with various downsampled resolutions. Here we followed the training protocol in the Histo-Cloud work^8^ using training patches randomly downsampled to 1, 2, 3, & 4 times smaller with respect to the native WSI resolution.

## Notes

### Competing Interest Statement

The authors have declared no competing interest.

